# Gut Microbiota Alterations Across REM Sleep Behavior Disorder and Parkinson’s Disease: A Machine Learning-Based Meta-Analysis

**DOI:** 10.1101/2025.04.09.647562

**Authors:** Muzaffer Arıkan

## Abstract

Recent studies have examined the relationship between rapid eye movement sleep behavior disorder (RBD), Parkinson’s disease (PD), and the gut microbiota, but no consensus exists on the shared and distinct gut microbiota changes. This study aimed to identify consistent and divergent gut microbiota changes across RBD and PD and to evaluate the performance of machine learning (ML) models in distinguishing PD, RBD, and healthy controls (HC). A meta-analysis of four gut microbiota studies involving PD, RBD, and HC groups was conducted, comprising a total of 973 samples (379 PD, 251 RBD, and 343 HC). ML models could differentiate PD from HC (cross study validation (CSV) AUC 0.61 ± 0.06) and RBD from HC (CSV AUC 0.58 ± 0.03). However, distinguishing between PD and RBD was ineffective (CSV AUC 0.51 ± 0.03). ML models distinguished PD and RBD from HC with weak to moderate predictive accuracy but failed to differentiate PD from RBD.

## Introduction

Parkinson’s disease (PD) is the second most common neurodegenerative disorder and the leading alpha-synucleinopathy, marked by the progressive loss of dopaminergic neurons in the substantia nigra pars compacta [1]. PD presents with a range of motor and non-motor symptoms, including gastrointestinal disturbances, loss of smell, and sleep disorders [2].

REM sleep behavior disorder (RBD) is a parasomnia characterized by recurrent “dream re-enactments” and loss of normal REM sleep muscle atonia. RBD can occur years before alpha-synucleinopathies and is therefore described as a prodromal phase for these disorders [3]. Over 80% of patients with idiopathic RBD progress to develop PD, dementia with Lewy bodies (DLB), or multiple system atrophy (MSA) within 14 years [3].

In recent years, several studies have investigated the relationship between RBD, PD and the gut microbiota [4–7]. However, there is no consensus on the overlapping and divergent gut microbiota changes in these two conditions, nor on the predictive performance of gut microbiota-based machine learning models to discriminate between them.

This study aimed to identify overlapping and divergent gut microbiota changes in RBD and PD with a comprehensive meta-analysis and to determine the performance of machine learning models based on existing gut microbiota data in distinguishing PD and RBD patients from healthy controls (HC).

## Methods

### Selected datasets

A PubMed search was conducted with no language restriction for publications from January 1, 2000, to March 1, 2025, using the search terms: (“Microbiome” OR “Microbiota”) AND (“REM” OR “sleep behavior disorder” OR “RBD”) AND (“Parkinson’s” OR “Parkinson”) NOT (Review[pt] OR “systematic review”[pt]). The study inclusion criteria were: (1) human studies involving PD, RBD and HC groups; (2) microbial DNA extraction from stool; and (3) amplicon sequencing of the 16S rRNA gene using next-generation sequencing.

### Data acquisition

For eligible studies that have deposited sequencing data in the NCBI SRA database, raw 16S rRNA gene sequences and subjects’ information were retrieved using the BioProject accession numbers provided in the publications.

### Bioinformatics

Raw paired-end 16S rRNA gene amplicon sequencing data were quality-filtered with Trimmomatic [8] and then merged with FLASH [9]. Taxonomic assignment and ASV abundance table generation were carried out using the DADA2 [10] and the SILVA database (v138.1) [11]. Amplicon sequence variants (ASVs) present in at least two samples were retained for downstream analyses, with samples rarefied to 2,000 reads prior to analysis. Data preparation, cleaning and diversity analyses were conducted using phyloseq [12] and microViz [13]. Differential abundance analysis and machine learning model construction were performed using SIAMCAT [14]. Specifically, LASSO algorithm was used to build the models and performances were tested on log normalized data. Model accuracies were evaluated through 10-fold cross-validation, repeated 10 times. Model performances were quantified by the area under the receiver operating characteristic curve (AUC).

Within-study cross-validations (CV) and cross-study validations (CSV) were conducted to test model performances across different datasets as previously described [15]. ML models were initially applied individually to each study using within-study CV to determine the ability of gut microbiota profiles to distinguish between HC, RBD, and PD samples. To assess the portability of these models, cross-study validation (CSV) was performed. In this approach, models built for each dataset were tested on all other datasets, and ML model accuracies were evaluated using AUC. Visualization of results was carried out using ggplot2.

## Results

Following the initial screening process, four studies were included in the meta-analysis, encompassing a total of 973 samples (378 PD, 249 RBD and 343 HC) across three countries. After data cleaning and filtering, a total of 970 samples remained for downstream analyses (**Table 1**).

**Table 1.**
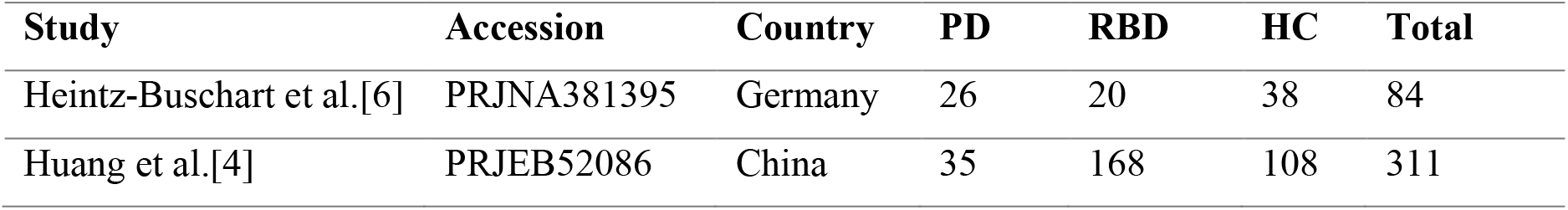
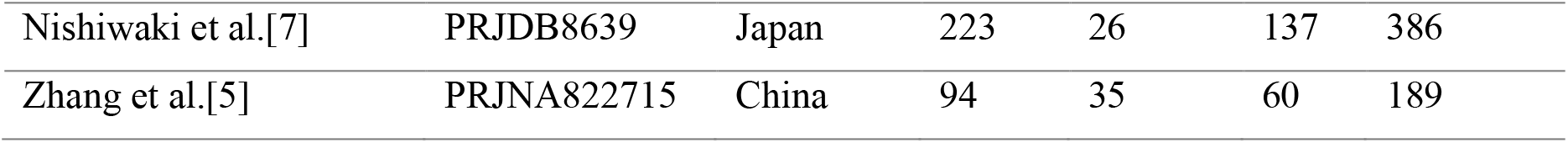
Overview of the studies used in this work. Samples refer to those used in the study after data filtration. HC, healthy controls; PD, Parkinson’s disease patients; RBD, Rapid eye movement sleep behavior disorder.

| Study | Accession | Country | PD | RBD | HC | Total |
| --- | --- | --- | --- | --- | --- | --- |
| Heintz-Buschart et al.[6] | PRJNA381395 | Germany | 26 | 20 | 38 | 84 |
| Huang et al.[4] | PRJEB52086 | China | 35 | 168 | 108 | 311 |
| Nishiwaki et al.[7] | PRJDB8639 | Japan | 223 | 26 | 137 | 386 |
| Zhang et al.[5] | PRJNA822715 | China | 94 | 35 | 60 | 189 |

### Gut microbiota diversity

Alpha diversity analysis revealed no significant differences among HC, RBD and PD groups in Heintz-Buschart et al. [6], Huang et al. [4] and Zhang et al. [5]. However, in Nishiwaki et al. [7], a significant difference was observed in the Shannon index (Kruskal-Wallis, p=0.0013) (**Figure 1a-b**).

**Figure 1.**
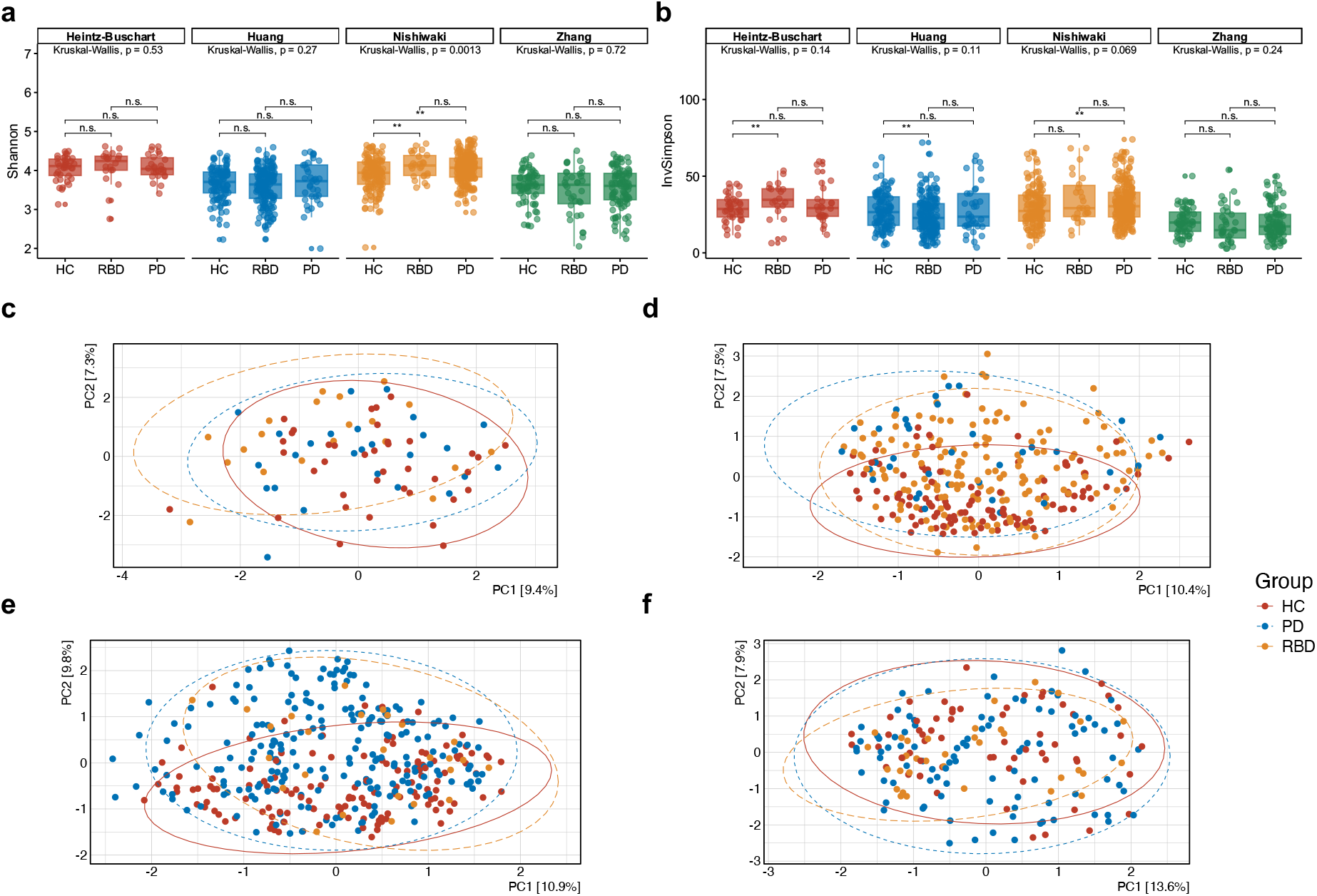
Alpha and beta diversity comparisons within individual studies. Alpha diversity comparisons based on the Shannon index (a) and inverse Simpson index (b) for samples grouped by disease status (HC, RBD, PD) within each study. Beta diversity comparisons between disease groups in Heintz-Buschart et al. [6] (c), Huang et al. [4] (d), Nishiwaki et al. [7] (e), and Zhang et al. [5] (f), based on principal coordinate analysis (PCoA) using Aitchison distance. Ellipses represent 95% confidence intervals, and colors indicate disease groups.

Beta diversity comparisons within individual studies showed significant differences between three disease groups in Huang et al. [4] (Aitchison, PERMANOVA, R^2^ = 0.012, p = 0.002), Nishiwaki et al. [7] (R^2^ = 0.012, p = 0.001), and Zhang et al. [5] (R^2^ = 0.013, *p* = 0.026), while no significant difference was observed in Heintz-Buschart et al. [6] (R^2^ = 0.029, p = 0.153) (**Figure 1c-f**).

Beta diversity analysis of pooled samples across studies indicated that microbial community structures varied by not only by disease group (**Figure 2a**) but also by study origin (**Figure 2b**). Multivariate analysis confirmed significant differences based on both the study of origin (PERMANOVA, R^2^ = 0.14, *p* = 0.001) and the disease group (PERMANOVA, R^2^ = 0.03, *p* = 0.001). Notably, the disease group explained only 3.6% of the variance, while the study of origin accounted for a much larger share at 14.4%, highlighting the noticeable cross-study variability commonly observed in microbiota meta-analyses.

**Figure 2.**
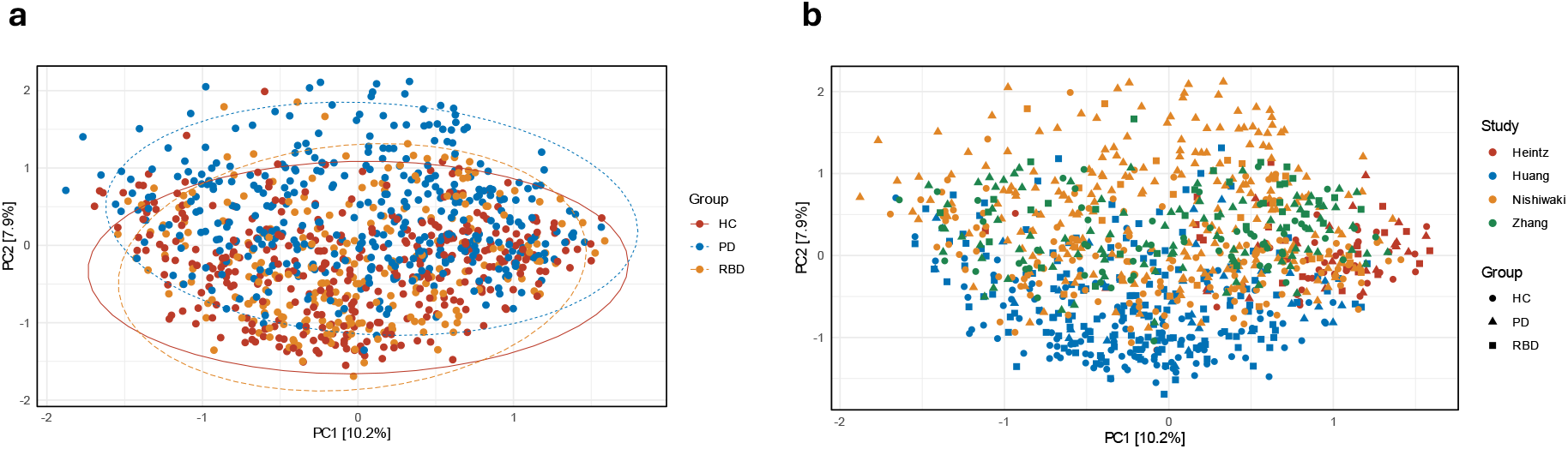
Beta diversity comparisons of pooled samples by disease group and study origin. (a) Principal coordinate analysis (PCoA) based on Aitchison distance, with ellipses indicating 95% confidence intervals and colors representing disease groups. (b) PCoA based on Aitchison distance, with colors indicating study of origin and shapes denoting disease groups.

### Gut microbiota features associated with RBD and PD

Differentially abundant genera identified in comparisons of RBD vs. HC, PD vs. HC and PD vs. RBD, which showed a strong association with the condition (single-feature AUROC > 0.6 or < 0.3) in at least one study, were extracted and visualized as heatmaps to display their generalized fold changes (**Figure 3a**).

**Figure 3.**
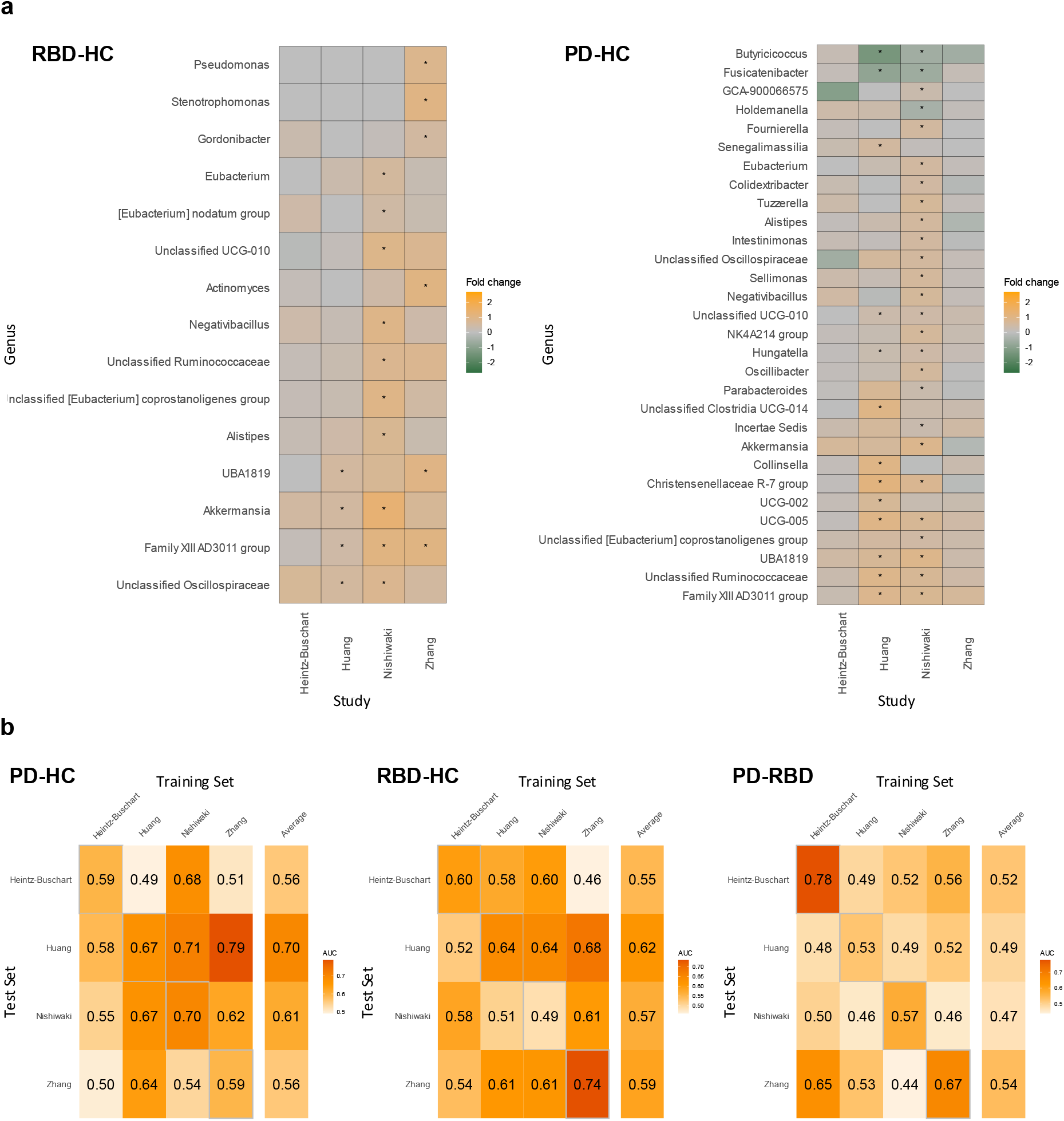
Genus-level differential abundance and association analysis for PD-HC and RBD-HC comparisons, along with cross-study portability of machine learning models. (a) Genera strongly associated with PD or RBD (single-feature AUROC > 0.6 or < 0.3) in at least one study were extracted and visualized as heatmaps to show their generalized fold changes. Differentially abundant genera (p < 0.05) in RBD-HC and PD-HC comparisons are marked with stars. (b) Classification accuracies of study-specific machine learning models were evaluated on other studies to assess model portability. Diagonal cells show cross-validation results (where test and training sets are the same), while off-diagonal cells represent study-to-study model transfer results. The average AUC for each study-specific model is displayed in the “Average” column of the heatmap.

In RBD vs. HC comparisons, an increased abundance of *Akkermansia, UBA1819, Family XIII AD3011*, and unclassified *Oscillospiraceae* was identified in at least two studies (**Figure 3a**). In PD vs. HC comparisons, a decreased abundance of *Butyricicoccus* and *Fusicatenibacter*, along with an increased abundance of unclassified *UCG-010, Hungatella, Christensenellaceae R-7* group, *UCG-005, UBA1819*, unclassified *Ruminococcaceae*, and *Family XIII AD3011*, were detected in at least two studies (**Figure 3a**). No differentially abundant taxa meeting the specified AUC range were detected in PD vs. RBD comparisons.

Shared gut microbiota changes associated with PD and RBD included an increased abundance of *UBA1819* and *Family XIII AD3011*. PD-specific changes included a decreased abundance of *Butyricicoccus* and *Fusicatenibacter*, with an increased abundance of unclassified *UCG-010, Hungatella, Christensenellaceae R-7* group, *UCG-005*, and unclassified *Ruminococcaceae*. RBD-specific changes were characterized by an increased abundance of *Akkermansia* and unclassified *Oscillospiraceae*.

### Machine learning model performances

For PD vs. HC classifications, Nishiwaki et al. [7] achieved the highest AUC (70%) in within-study CV, while Heintz-Buschart et al. [6] and Zhang et al.[5] had the lowest (59%). For RBD vs. HC, Zhang et al. [5] attained the highest AUC (74%), and Nishiwaki et al. [7] the lowest (49%) (**Figure 3b**). Models built on Huang et al. [4] and Zhang et al. [5] datasets performed better when applied to each other’s dataset. In PD vs. RBD classifications, Heintz-Buschart et al. [6] reached the highest AUC (78%) in within-study CV, while Huang et al. [4] had the lowest (53%) (**Figure 3b**). ML models could differentiate PD from HC with a within-study average CV AUC of 0.63 ± 0.05, RBD from HC with a within-study average CV AUC of 0.61 ± 0.10, and PD from RBD with a within-study average CV AUC of 0.63 ± 0.11.

Compared to the performance estimated through within-study CV, CSV performances were not significantly different in any of the group comparisons (two samples Welch t-test: PD vs. HC t = −0.69, df = 5.85, p = 0.52; RBD vs. HC t = 0.65, df = 3.49, p = 0.56; PD vs. RBD t = 2.28, df = 3.46, p = 0.09). There was no correlation between AUC in the within-study CV and average CSV AUC for ML models (Pearson, r = 0.47, p = 0.24). Average CSV AUCs obtained for PD vs. HC models were not significantly different from RBD vs HC models (two samples Welch t-test: t = 0.69, df = 4.17, p-value = 0.53; average AUC PD vs. HC 60.8% ± 6.6, average AUC RBD vs. HC 58.3% ± 2.98). However, average CSV AUCs obtained for both PD vs. HC (two samples Welch t-test: t = 2.80, df = 4.27, p-value = 0.04; average AUC PD vs. HC 60.8% ± 6.6, average AUC PD vs. RBD 50.5% ± 3.11) and RBD vs. HC models (two samples Welch t-test: t = 3.59, df = 5.99, p = 0.01; average AUC RBD vs. HC 58.3% ± 2.98, average AUC PD vs. RBD 50.5% ± 3.11) were higher than PD vs. RBD models. ML models could differentiate PD from HC (average CSV AUC 0.61 ± 0.06) and RBD from HC (average CSV AUC 0.58 ± 0.03). However, distinguishing between PD and RBD was ineffective (average CSV AUC 0.51 ± 0.03).

## Discussion

In this study, a ML-based meta-analysis of gut microbiota studies was conducted to explore alterations across RBD and PD, as well as to assess the predictive performance of ML models in distinguishing PD, RBD and HC. The analysis identified shared and distinct microbiota changes across multiple studies, with varying classification accuracy for microbiota profile-based ML models.

Microbial diversity analyses within individual studies revealed partially overlapping patterns. Alpha diversity did not differ significantly between HC, RBD and PD groups in three of the studies, with the exception of Nishiwaki et al. [7]. In contrast, beta diversity differed significantly in all but one study, Heintz-Buschart et al. [6]. When samples were pooled for beta diversity analysis, study origin emerged as a major driver of variation in gut microbial composition along with disease status, reflecting the frequently observed heterogeneity in microbiome research, as highlighted in previous meta-analyses [15].

In RBD vs. HC comparisons, an increased abundance of *Akkermansia, UBA1819, Family XIII AD3011*, and unclassified *Oscillospiraceae* was identified in at least two studies. For PD vs. HC comparisons, a decreased abundance of *Butyricicoccus* and *Fusicatenibacter*, along with an increased abundance of unclassified *UCG-010, Hungatella, Christensenellaceae R-7* group, *UCG-005, UBA1819*, unclassified *Ruminococcaceae*, and *Family XIII AD3011*, was detected in at least two studies. While differential abundance findings generally aligned with original studies, no taxa were consistently identified across all studies, likely due to variability and low statistical power from small sample sizes in the studies.

ML models could differentiate PD and RBD from HC with weak to moderate predictive performance in both within-study CV and CSV evaluations. The average CSV AUCs for both PD vs. HC and RBD vs. HC models were higher than those for PD vs. RBD models. Although not statistically significant, there was a notably lower average CSV AUC for ML models in distinguishing PD from RBD compared to within-study CV evaluations. This suggests that while ML models could differentiate PD from RBD with weak to moderate predictive performance in within-study CV evaluations, they failed in CSV evaluations, indicating that these models were more study-specific with low cross-study portability. Moreover, no correlation was observed between within-study CV and the cross-study portability of the models.

This study has several limitations. First, due to the lack of available metadata, the identified microbiota features and ML models might be potentially confounded by other donor characteristics, such as age, sex, or medication usage. Second, the primary goal of the study was not a comprehensive evaluation of all parameter optimizations and ML algorithms, which means differences in predictive performance may exist based on different algorithms or parameters.

In conclusion, this study identifies both overlapping and distinct alterations in the human gut microbiota associated with PD and RBD while assessing the potential of ML models to differentiate between PD, RBD, and HC. The findings underscore the challenges in the portability of gut microbiota-based ML models and their limitations in distinguishing RBD from PD. More comprehensive multi-omics and multi-site approaches may be required to achieve better discrimination between PD and RBD, improving predictive accuracy and early-stage disease differentiation. Future research should consider longitudinal studies to track how gut microbiota changes over time in PD and RBD, which could provide valuable insights into disease progression. Additionally, expanding the sample size and incorporating a broader range of clinical variables may further refine the accuracy of machine learning models, offering more robust tools for early diagnosis and differentiation.

## Acknowledgement

The author thanks all authors who made their datasets available.

## Data availability

The raw sequence data analyzed in this study are available under accession numbers: PRJNA381395, PRJEB52086, PRJDB8639 and PRJNA822715.

## Competing interests

The author declares no competing interests.

## Notes

### Competing Interest Statement

The authors have declared no competing interest.

